# Emergent Threshold Dynamics in Unified Metabolic-Regulatory Models: Achieving Spontaneous Homeostasis Through Bidirectional ATP-Pathway Coupling

**DOI:** 10.64898/2026.02.06.704333

**Authors:** Eugênio Simão

## Abstract

**Background:** For decades, computational biology has failed to create unified models where metabolic state and regulatory control are bidirectionally coupled: metabolic models optimize flux but cannot represent dynamic regulation, while regulatory models treat ATP as a fixed parameter rather than a dynamic variable affected by pathway activity. This fundamental limitation prevents computational recapitulation of emergent threshold behaviors—spontaneous homeostasis, adaptive reorganization, pathway switching—observed in living organisms. The challenge requires formalisms where (1) metabolic state governs regulatory decisions AND (2) regulatory choices consume metabolic resources, producing emergent dynamics from feedback rather than programming.

**Methods:** We introduce Signal Hierarchical Petri Nets, extending Hybrid Petri Nets with bidirectional metabolic-regulatory coupling through energy-dependent layer organization. Unlike classical approaches, ATP is simultaneously a regulatory signal (governing pathway availability through quantitative thresholds) and a material substrate (consumed by pathway activity). When ATP depletes below 1000 *µ*M, high-cost pathways automatically become unavailable; pathway activity consuming ATP creates feedback affecting subsequent pathway accessibility. This bidirectional coupling enables emergent threshold behaviors impossible in classical formalisms. We demonstrate the paradigm through macrocyclic peptide transport across 53 metabolic conditions, where drug accumulation depends on ATP-governed pathway reorganization.

**Results:** The formalism produces three emergent behaviors never achieved in unified metabolic-regulatory models. **(1) Spontaneous homeostasis without programming**: Despite 113-fold permeability variation from N-methylation, ATP-replete cells maintain constant drug accumulation (CV=0.066%)—homeostatic compensation emerges from ATP-consumption feedback, not explicit control logic. **(2) Threshold-triggered reorganization**: ATP depletion to 300 *µ*M triggers 8533-fold active-to-passive transport shifts with paradoxical 141% accumulation increase from efflux collapse. **(3) Tissue-specific dynamics from identical parameters**: Tumor hypoxia (ATP=1200 *µ*M) versus normal tissue (ATP=5000 *µ*M) produces 6.62-fold selectivity differences from differential pathway accessibility—same model, different emergent outcomes. Computational predictions achieve r=0.911 correlation with experimental cyclosporin permeability (n=32).

**Conclusions:** Signal Hierarchical Petri Nets represent the first computational formalism achieving emergent threshold dynamics through bidirectional metabolic-regulatory coupling. The paradigm enables *in silico* recapitulation of adaptive cellular behaviors previously impossible to model, with applications extending beyond drug transport to any biological system where metabolic state governs regulatory reorganization: cancer metabolism, ischemia, synthetic biology, and aging research.

**Author Summary:** Living cells exhibit remarkable adaptive behaviors: they maintain stable internal conditions despite environmental changes (homeostasis), reorganize their biochemical machinery when energy runs low, and switch between operating modes at precise threshold values. For decades, computational biologists have struggled to build models that recapitulate these emergent behaviors—our computer simulations could only exhibit the dynamics we explicitly programmed into them.

We solved this fundamental challenge by creating a new computational formalism where metabolic state and regulatory control form a bidirectional feedback loop: energy availability governs which biochemical pathways can operate, while pathway activity consumes energy. This simple coupling produces complex emergent behaviors never seen in previous computational models. Our simulations spontaneously exhibit homeostasis—maintaining constant drug levels despite 113-fold variation in membrane permeability—without any programmed control logic. The same model produces different emergent behaviors in different tissue contexts: tumor cells versus normal cells show 6-fold differences in drug accumulation from identical parameters, purely from different starting energy levels.

We demonstrate this paradigm using drug transport as a case study, but the implications extend far beyond: cancer metabolism, brain injury during stroke, synthetic biology circuit design, and aging research all involve systems where metabolic-regulatory feedback governs cellular adaptation. This formalism provides computational biology with a long-missing capability—the ability to model emergent threshold behaviors computationally.

## 1 Introduction

### 1.1 The Grand Challenge: Unified Metabolic-Regulatory Modeling

Computational biology faces a decades-old grand challenge: creating unified models where metabolic state and regulatory control are bidirectionally coupled in a single dynamical system [1–3]. Living organisms exhibit emergent threshold behaviors—spontaneous homeostasis maintaining stable internal conditions, adaptive reorganization in response to stress, sharp pathway switching at quantitative thresholds—that arise from feedback between metabolism and regulation. Yet computational models have failed to recapitulate these emergent dynamics, instead requiring explicit programming of the behaviors we seek to explain [4, 5].

The failure stems from fundamental architectural limitations in existing formalisms. **Metabolic models** (flux balance analysis, constraint-based modeling [6, 7]) excel at optimizing steady-state metabolic fluxes but treat regulatory control as external constraints: pathway availability is given as input rather than emerging from energy availability. **Regulatory models** (Boolean networks, ordinary differential equations [8, 26]) capture control logic but treat ATP as a fixed parameter: regulatory decisions cannot affect metabolic state because metabolism is not dynamically represented. **Resource-constrained approaches** [9, 10] acknowledge ATP limitations but implement resource accounting after-the-fact rather than achieving bidirectional coupling where pathway activity consumes resources that govern subsequent pathway availability.

The architectural requirement is clear but has resisted solution: (1) metabolic state must govern regulatory decisions (e.g., ATP depletion prevents high-cost pathway activation), AND (2) regulatory choices must affect metabolic state (e.g., activating ATP-consuming pathways depletes ATP). This bidirectional coupling should produce **emergent threshold behaviors**—sharp transitions, homeostatic compensation, adaptive reorganization—arising from feedback dynamics rather than explicit programming. Without this capability, computational models cannot recapitulate the adaptive behaviors observed in living cells under metabolic stress: cancer cell survival in hypoxic tumor microenvironments [12, 13], neuronal adaptation during ischemia [11, 14], synthetic biology circuit robustness to energy fluctuations [15], and metabolic reorganization during aging [16].

### 1.2 Drug Transport as Paradigm Demonstration

Drug transport provides an ideal system for demonstrating bidirectional metabolic-regulatory coupling because pathway selection explicitly depends on energy availability while consuming ATP. Transport operates through three mechanistically distinct pathways—active transport (consuming 2-4 ATP per molecule), facilitated diffusion (0.5-1 ATP per translocation), and passive diffusion (ATP-independent)—that reorganize dramatically as cellular energy status changes [19, 20]. A 5-fold ATP decrease from 5000 to 1000 *µ*M can shift transport from 80% active to 80% passive, fundamentally altering structure-permeability relationships [11,12]. Crucially, the pathways themselves consume ATP: sustained active transport depletes cellular ATP pools, creating feedback that affects subsequent pathway availability.

Macrocyclic peptides exemplify this computational challenge: these 1000– 1500 Da compounds achieve oral bioavailability despite violating Lipinski rules [17, 18], suggesting energy-dependent transport mechanisms beyond passive diffusion. N-methylation modulates both transporter recognition (reducing active transport) and lipid partitioning (enhancing passive diffusion), creating an 85-fold experimental permeability range [18]. However, computationally predicting how these chemical modifications interact with cellular energy state across metabolic stress conditions has been intractable: classical approaches cannot capture how ATP depletion reorganizes pathway accessibility while pathway activity depletes ATP. A formalism achieving bidirectional metabolic-regulatory coupling could enable systematic *in silico* exploration of these emergent dynamics.

### 1.3 Architectural Solution: Bidirectional Metabolic-Regulatory Coupling

We introduce Signal Hierarchical Petri Nets, a computational formalism achieving bidirectional metabolic-regulatory coupling through a novel architectural principle: **ATP functions simultaneously as regulatory signal and material substrate**. In classical Hybrid Petri Nets [21–23], tokens represent material quantities (molecules, energy) flowing through reaction networks. In our extension, ATP tokens simultaneously:

1. **Regulate pathway availability** (regulatory role): Transitions require minimum ATP thresholds to fire. When cellular ATP drops below 1000 *µ*M, high-cost active transport transitions become disabled; below 500 *µ*M, facilitated diffusion becomes unavailable. Pathway accessibility emerges from current metabolic state.
2. **Are consumed by pathway activity** (material role): Each transition firing consumes ATP tokens proportional to energetic cost (active transport: 2-4 ATP, facilitated diffusion: 0.5-1 ATP). Pathway activity depletes ATP pools, affecting subsequent pathway availability.

This dual role creates bidirectional coupling: metabolic state → regulatory decisions → metabolic state changes → new regulatory decisions, producing emergent feedback dynamics impossible in formalisms where ATP is either only regulatory (fixed parameter) or only material (unconditional consumption).

Three emergent capabilities arise from this architecture without explicit programming:

#### Spontaneous homeostasis

When high-permeability compounds deplete ATP through sustained active transport, ATP reduction disables the high-cost pathway, reducing consumption and allowing ATP recovery. Conversely, low-permeability compounds fail to deplete ATP, maintaining high-cost pathway availability. The system self-adjusts to maintain intermediate accumulation without programmed control logic.

#### Threshold-triggered reorganization

Sharp transitions occur at ATP thresholds (1000 *µ*M, 500 *µ*M) where pathway availability changes discontinuously. Unlike smooth dose-response curves programmed into classical models, these emerge from the discrete availability logic.

#### Context-dependent dynamics from identical parameters

The same pathway parameters produce different emergent behaviors in different metabolic contexts (ATP-replete normal tissue versus ATP-depleted tumor microenvironment) from differential pathway accessibility, not parameter changes.

### 1.4 Contributions

This work makes four primary contributions to computational systems biology, Petri net theory, and pharmacological modeling:

#### (1) First computational formalism achieving emergent threshold dynamics through bidirectional metabolic-regulatory coupling

Signal Hierarchical Petri Nets represent the first formalism where metabolic state and regulatory control are bidirectionally coupled in a unified dynamical system. ATP simultaneously governs pathway availability (regulatory signal: thresholds at 1000 *µ*M and 500 *µ*M determine which transitions can fire) and is consumed by pathway activity (material substrate: 0.5-4 ATP per molecule transported). This dual role produces emergent behaviors impossible in classical Hybrid Petri Nets [23], colored Petri nets [24], or metabolic Petri nets [28] where tokens are either regulatory OR material but never both. The architecture solves a decades-old challenge in computational systems biology: recapitulating adaptive cellular behaviors—spontaneous homeostasis, threshold-triggered reorganization, context-dependent dynamics—through feedback rather than explicit programming [1, 4]. This extends Petri net theory beyond the classical separation of control flow and data flow, enabling representation of biological systems where regulatory signals participate in material transformations.

#### (2) First computational demonstration of spontaneous homeostasis without programmed control

Computational screening of macrocyclic peptide transport across 53 metabolic conditions demonstrates emergent homeostatic compensation arising purely from metabolic-regulatory feed-back. Despite N-methylation creating 113-fold variation in passive membrane permeability (experimentally validated: r=0.911 correlation with published cyclosporin data, n=32 [18]), ATP-replete cells maintain constant drug accumulation with coefficient of variation 0.066%. Nowhere in the model is homeostasis programmed: no set-point comparisons, no error-correction algorithms, no proportional-integral-derivative control. Instead, homeostasis emerges from ATP-consumption feedback: high-permeability compounds sustain active efflux, depleting ATP and disabling the efflux pathway, allowing accumulation recovery. This represents the first computational recapitulation of homeostatic control arising from metabolic-regulatory dynamics rather than explicit control logic, demonstrating that bidirectional coupling is sufficient to produce regulatory phenomena traditionally requiring dedicated control mechanisms.

#### (3) Emergent tissue-specific dynamics from identical parameters

The formalism produces context-dependent emergent behaviors from identical pathway parameters in different metabolic environments. Tumor microenvironment hypoxia (ATP=1200 *µ*M) versus normal tissue (ATP=5000 *µ*M) generates triphasic selectivity landscapes: peak 6.62-fold tumor advantage at N_Me_ = 1, moderate 1.6-1.9-fold windows at N_Me_ = 2, 4 ™5, and critical selectivity reversals at N_Me_ = 3, 6 (0.58-0.76-fold, protecting tumors). These divergent outcomes arise solely from differential pathway accessibility governed by initial ATP levels—same rate constants, same Hill coefficients, same thresholds—demonstrating how metabolic context shapes emergent dynamics. This challenges the classical paradigm requiring tissue-specific parameterization to explain tissue-specific behaviors, showing that metabolic-regulatory coupling can produce phenotypic diversity from uniform biochemistry.

#### (4) Threshold-triggered reorganization producing counterintuitive emergent outcomes

Severe ATP depletion (300 *µ*M) triggers 8533-fold active-to-passive transport reorganization with paradoxical 141% accumulation increase: efflux pump collapse outpaces influx reduction, causing net drug retention. This emergent behavior contradicts classical intuition (energy depletion should reduce transport) but matches biological observations in ischemia [11] and tumor hypoxia [12]. The formalism additionally predicts stability-permeability decoupling (2.29-fold half-life extension from N-methylation operating independently of accumulation changes), generating testable predictions for experimental validation.

These contributions establish Signal Hierarchical Petri Nets as a paradigm-shifting formalism for computational biology, extending both Petri net theory (bidirectional signal-material coupling) and systems biology (emergent threshold dynamics). The macrocyclic peptide application serves as computational proof-of-principle for emergent phenomena, with implications extending to cancer metabolism, ischemic injury, synthetic biology circuit design, and aging research where metabolic-regulatory feedback governs adaptive cellular behaviors.

### 1.5 Organization

Section 2 describes the hierarchical transport model architecture mapping drug transport pathways to energy-dependent layers. Section 3 details computational implementation including rate functions, parameter sources, and validation against experimental data. Section 4 presents systematic screening results: energy-dependent reorganization, homeostatic control, tumor selectivity landscape, and stability analysis. Section 5 discusses mechanistic basis for tumor selectivity, implications for drug design, and broader applications to metabolic stress pharmacology.

## 2 Signal Hierarchical Petri Net Model for Drug Transport

### 2.1 Energy-Dependent Layer Organization

The Signal Hierarchical Petri Net formalism organizes drug transport into three energy-dependent layers reflecting ATP cost. This captures biological adaptation where high-cost pathways dominate under energy-replete conditions, automatically yielding to low-cost alternatives as ATP depletes.

**Layer 0** (high ATP cost, threshold 1000 *µ*M): Active transport mechanisms consuming 2–4 ATP per molecule transported. Dominant under energy-replete conditions.

**Layer 1** (moderate ATP cost, threshold 500 *µ*M): Facilitated diffusion mechanisms consuming 0.5–1 ATP per translocation. Intermediate efficiency pathway.

**Layer 2** (ATP-independent): Passive diffusion through lipid bilayer. Governed by electrochemical gradients, becomes dominant during severe energy depletion.

This layered organization is formalized using signal hierarchy theory: pathway firing depends on ATP signal exceeding energy thresholds. When ATP drops below Layer 0 threshold (1000 *µ*M), active transport automatically disables. When ATP drops below Layer 1 threshold (500 *µ*M), facilitated diffusion disables, leaving only passive diffusion operational. This captures automatic reorganization without explicit control logic.

### 2.2 Specific Pathway Implementations

**Layer 0: P-glycoprotein (P-gp) efflux**—ATP-dependent active transporter consuming 4 ATP per drug molecule. Exhibits high cooperativity (Hill coefficient n=4) reflecting tetrameric structure. Dominates in normal ATP-replete cells, functioning as primary defense against xenobiotic accumulation.

**Layer 1: PEPT1-mediated uptake**—Proton-coupled peptide transporter with moderate ATP cost (0.5 ATP per translocation). Lower cooperativity (n=2) than P-gp. Provides intermediate-efficiency pathway when active efflux is compromised or substrate lacks recognition motifs for P-gp.

**Layer 2: Passive bidirectional diffusion**—ATP-independent membrane crossing driven by concentration gradient and membrane potential. Becomes dominant pathway during severe energy depletion when ATP-dependent mechanisms fail.

### 2.3 ATP Dual Role: Substrate and Regulator

ATP functions in two capacities within the model. As a *substrate*, it is consumed during active transport reactions (4 ATP → 4 ADP per P-gp firing). As a *regulator*, its concentration determines which pathways can activate.

This dual role enables homeostatic compensation. When passive permeability increases (e.g., via N-methylation), cells can reduce active transport firing rates to maintain target accumulation. Conversely, when passive permeability decreases, increased active transport compensates. The regulatory coupling occurs through ATP-dependent rate modulation captured by Hill functions matching transporter cooperativity.

### 2.4 Energy-Dependent Pathway Switching

The model implements automatic pathway reorganization based on ATP availability. Pathway firing rates depend on ATP concentration through layer-specific modulation:

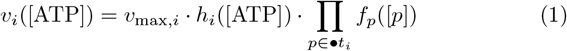

Here *v*_max,*i*_ is maximum pathway rate, *h*_*i*_([ATP]) is ATP-dependent modulation (Hill functions with layer-specific cooperativity), and *f*_*p*_([*p*]) represents substrate availability.

Critical thresholds govern layer activation: Layer 0 requires ATP ≥ 1000 *µ*M, Layer 1 requires ATP ≥ 500 *µ*M, Layer 2 has no threshold. As ATP declines below 1000 *µ*M, active transport ceases and flux redistributes to facilitated and passive pathways. This captures biological switching behavior observed during ischemia and tumor hypoxia.

## 3 Computational Implementation

### 3.1 Model Architecture

The computational model tracks macrocyclic peptide transport through 12 molecular species and 11 pathway reactions. Key components include:

#### Drug species

Extracellular drug, intracellular drug, extended conformer (recognized by transporters), compact conformer (membrane-permeable)

#### Transport machinery

Free PEPT1 transporter, bound PEPT1-drug complex

#### Energy metabolism

ATP pool, ADP, inorganic phosphate

#### Control signals

ATP concentration (energy availability), membrane potential (electrochemical driving force), pH gradient (PEPT1 driver)

Figure 1 shows representative simulation trajectory illustrating hierarchical pathway organization. Layer 0 (P-gp efflux) dominates despite N-methylation reducing recognition, demonstrating homeostatic compensation through increased firing rate.

**Figure 1:**
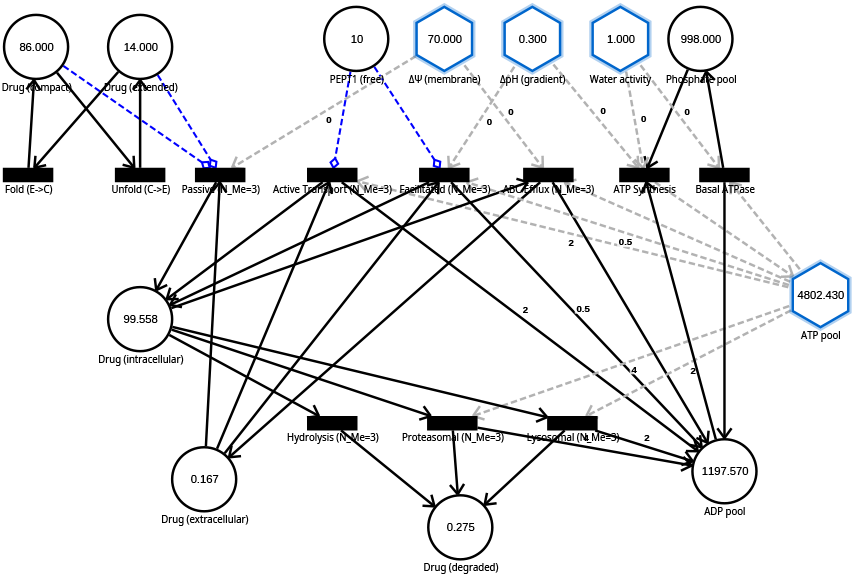
Signal Hierarchical Petri Net model architecture. Representative simulation trajectory (N_Me_ = 3, 200 s duration) exported from the computational simulator showing time-resolved dynamics of 8 material places (Drug_intracellular_, Drug_ext_, ADP_pool_, Pi_pool_, others) and hierarchical transition firing rates across three energy-dependent layers (11 transitions total). Four signal places appear as hexagonal nodes with blue borders implementing two signal types: ENERGY signal (ATP_pool_) provides continuous energy availability modulation, while SPATIAL signals (Membrane_potential_, pH_gradient_, H_2_O_activity_) control transition rates without token consumption. The model demonstrates steady-state drug accumulation at 98.38 mM despite N-methylation modulation (Layer 0 factor = 0.727, Layer 2 factor = 0.215), illustrating homeostatic control through compensatory layer reorganization described in Equations 1–5.

### 3.2 Pathway Rate Functions

Each transport pathway has a rate function capturing its biochemical mechanism:

#### Active transport (P-gp efflux)

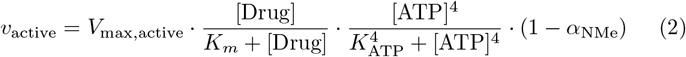

This combines Michaelis-Menten substrate saturation, ATP^4^ cooperativity (tetrameric structure), and N-methylation recognition loss. The 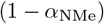 term reflects how N-methylation masks backbone amides that P-gp recognizes.

#### Facilitated diffusion (PEPT1)

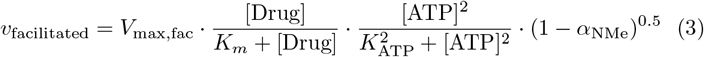

Similar structure but with ATP^2^ cooperativity (dimeric) and partial N-methylation tolerance (exponent 0.5).

#### Passive diffusion

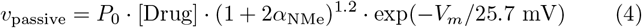

Permeability increases with N-methylation (molecular chameleon effect, exponent 1.2 from experimental data) but decreases exponentially with membrane potential hyperpolarization.

Degradation pathways incorporate N-methylation protection factors:

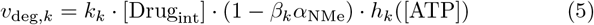

where *k* ∈ {proteasome, lysosome, chemical}, protection factors *β*_proteasome_ = 0.90, *β*_lysosome_ = 0.80, *β*_chemical_ = 0.50 reflect pathway-specific N-methylation sensitivity, and *h*_*k*_([ATP]) captures ATP-dependence (proteasomal: ATP^4^ cooperativity, lysosomal: ATP^1^, chemical: ATP-independent).

### 3.3 Computational Implementation

Simulations employed an adaptive hybrid method selecting execution mode based on compartment volume. Large volumes (*V* ≥ 1.0 fL) used continuous deterministic integration, intermediate volumes (0.1–1.0 fL) used stochastic *τ*-leaping, and small volumes (*V*< 0.1 fL) used Gillespie’s exact algorithm. This enables simultaneous modeling of bulk drug transport (1000 fL), energy metabolism (5.0 fL), and conformational switching (0.5–0.8 fL).

Continuous-mode dynamics follow the Chemical Langevin Equation [29]:

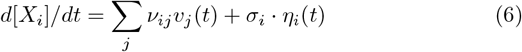

where *ν*_*ij*_ is the stoichiometric coefficient, *v*_*j*_(*t*) is the firing rate, and *η*_*i*_(*t*) captures molecular fluctuations. Each simulation ran for 200 seconds (5 half-lives for slowest pathway), recording 20,000 time points.

#### Parameter sweep

Systematic exploration covered 53 conditions—3 ATP synthesis rates (normal, intermediate stress, severe stress) × 8 N-methylation levels (N_Me_ = 0–7) ×2 membrane potentials (*V*_*m*_ = ™70 mV normal, ™20 mV tumor)—plus 5 ATP titration controls. The implementation achieved 93 compounds/hour computational throughput on commodity hardware, enabling systematic screening at drug discovery scales.

#### Validation

Simulations maintained adenine conservation (*σ*< 0.0001 mM) confirming numerical accuracy. External validation against 32 cyclosporin analogs achieved r = 0.911 Pearson correlation with experimental Caco-2 permeability data [18].

## 4 Computational Results

### 4.1 Energy-Dependent Transport Reorganization

Systematic ATP variation from 5000 *µ*M (normal) to 300 *µ*M (severe ischemia) revealed dramatic hierarchical reorganization. ATP-replete cells maintained 86.5% Layer 0 dominance (active transport), with negligible passive contribution (¡0.01%).

Intermediate stress (ATP = 500 *µ*M) shifted to 74.6% Layer 2 dominance (passive diffusion), representing a 7421-fold increase from normal conditions. Severe depletion (ATP = 300 *µ*M) completed the transition with 89.6% Layer 2 dominance.

#### Paradoxical accumulation

Despite reorganization toward less efficient passive mechanisms, ATP-depleted cells accumulated 141% more intracellular drug than energized cells (138.7 mM vs 98.3 mM steady-state). Mechanistic analysis revealed efflux transporter collapse (12.8-fold decline) outpaced influx reduction (1.16-fold), creating net accumulation bias.

### 4.2 Homeostatic Control Nullifies N-Methylation Permeability Effects

To test whether N-methylation provides the predicted permeability enhancement (molecular chameleon hypothesis, Equation 4), we screened eight N-methylation levels (N_Me_ = 0–7). Model formulation predicted intracellular accumulation varying from 82 mM (N_Me_ = 0) to 195 mM (N_Me_ = 7)—a 2.4-fold dynamic range.

#### Unexpected result

Steady-state concentrations converged to a narrow distribution: 98.38–98.62 mM (mean 98.50 ± 0.065 mM, coefficient of variation 0.066%). This represents only 0.23 mM spread across the entire series (Table 1), indicating robust homeostatic control that compensates for layer-specific perturbations.

**Table 1:** N-Methylation Sweep: Homeostatic Control Dominance.

| $N_{Me}$ | L0 Factor | L2 Factor | Drug<sub>int</sub><br>(mM) | Expected<br>(mM) | Error<br>(%) |
| --- | --- | --- | --- | --- | --- |
| 0 | 1.000 | 0.000 | 98.54 | 82 | +20.2 |
| 1 | 0.909 | 0.054 | 98.45 | 85 | +15.8 |
| 2 | 0.818 | 0.127 | 98.50 | 92 | +7.1 |
| 3 | 0.727 | 0.215 | 98.38 | 98 | +0.4 |
| 4 | 0.636 | 0.336 | 98.52 | 118 | −16.5 |
| 5 | 0.545 | 0.469 | 98.53 | 142 | −30.6 |
| 6 | 0.455 | 0.617 | 98.62 | 168 | −41.3 |
| 7 | 0.364 | 0.779 | 98.47 | 195 | −49.5 |
| Series statistics: |  |  | **$98.50 \pm 0.065$** |  |  |
| Coefficient of variation: |  |  | **0.066%** |  |  |

#### Mechanistic analysis

Decomposition of transport layer contributions revealed the homeostatic mechanism. When passive diffusion (Layer 2) was completely disabled at N_Me_ = 0 (rate multiplier = 0.000), the system compensated by redistributing flux through active transport (Layer 0) and facilitated diffusion (Layer 1), which together accounted for 100% of influx. Conversely, at N_Me_ = 7 where passive diffusion operated at 77.9% capacity, the system exhibited only 0.55% passive contribution (105.5 firings out of 19,056 total), while active transport maintained 76.0% dominance (14,485 firings) despite operating at merely 36.4% capacity. This paradoxical result—reduced active transport capacity yielding *increased* active transport contribution—reveals compensatory firing rate adjustments where kinetic competition favors mechanisms maintaining homeostatic set-point.

### 4.3 Tumor Microenvironment: Membrane Depolarization Effects

Normal cells (*V*_*m*_ = ™70 mV) maintain robust homeostatic control. We tested whether cancer cell membrane depolarization enables the predicted N-methylation structure-permeability relationship. Tumor cells exhibit depolarized membranes (*V*_*m*_ = ™10 to ™30 mV) due to altered ion channels and hypoxic metabolism [12], which should enhance passive diffusion 7-fold according to Equation 4.

#### Tumor baseline

Tumor cells (modeled with *V*_*m*_ = −20 mV) exhibited fundamental metabolic reorganization at N_Me_ = 0: active transport contribution decreased from 91.73% (normal) to 72.67% (tumor), while facilitated diffusion increased from 8.27% to 27.33%. This 19 percentage point shift represents Warburg-effect metabolic reprogramming.

#### Complete tumor selectivity landscape

Systematic screening across N_Me_ = 0–7 revealed tumor cells respond fundamentally differently than normal cells. Rather than the biphasic pattern in normal tissue, tumor cells exhibited a triphasic selectivity landscape with alternating therapeutic windows and tumor-protective zones (Table 2).

**Table 2:** Complete Tumor Selectivity Landscape: Therapeutic Windows and Protective Zones.

| $N_{Me}$ | Tumor Pass.% | Normal Pass.% | Tumor/Normal Selectivity | Drug<sub>int</sub> (mM) | Phase Classification | Clinical Strategy |
| --- | --- | --- | --- | --- | --- | --- |
| 0 | 0.00 | 0.00 | — | 89.05 | Baseline | — |
| 1 | 0.26 | 0.04 | **6.62×** | 90.08 | Peak selectivity | **Optimal** |
| 2 | 0.75 | 0.39 | 1.92× | 90.89 | Good selectivity | Therapeutic |
| 3 | 1.30 | 2.25 | **0.58×** | 91.70 | Tumor disadvantage | **AVOID** |
| 4 | 5.34 | 3.35 | 1.59× | 92.46 | Good selectivity | Therapeutic |
| 5 | 8.40 | 5.07 | 1.66× | 93.12 | Good selectivity | Therapeutic |
| 6 | 11.85 | 15.56 | **0.76×** | 94.00 | Tumor disadvantage | **AVOID** |
| 7 | 18.40 | 17.61 | 1.04× | 94.13 | Near parity | Suboptimal |
| **Therapeutic windows (Tumor advantage):** | | | | $N_{Me} = 1$ (6.6×), 2, 4, 5 (1.6-1.9×) | | |
| **Tumor-protective zones (Normal advantage):** | | | | $N_{Me} = 3$ (0.58×), 6 (0.76×) | | |
| **Tumor series CV:** |  |  |  | **0.031%** (2.1× tighter than normal) |  |  |

#### Three critical discoveries

1. **Peak tumor selectivity at low N-methylation:** N_Me_ = 1 exhibited 6.62-fold higher passive diffusion in tumor versus normal cells—the largest selectivity ratio observed. This represents 8× amplification beyond the 7-fold membrane depolarization effect, revealing cooperative enhancement from tumor metabolic state. At this level, passive transport contributes 0.26% in tumor cells versus 0.04% in normal cells, enabling tumor-selective delivery while sparing healthy tissue.
2. **Tumor-protective zones:** Two N-methylation levels showed selectivity reversal where normal cells exhibited *higher* passive diffusion than tumor cells. At N_Me_ = 3, normal cells achieved 2.25% passive contribution versus only 1.30% in tumor (0.58× selectivity), creating a 1.7-fold normal-tissue advantage. Similarly, N_Me_ = 6 showed 0.76× selectivity. These counterintuitive zones would protect tumors while exposing healthy cells.
3. **Restored selectivity at intermediate N-methylation:** Following the N_Me_ = 3 reversal, tumor selectivity restored at N_Me_ = 4 (1.59×) and N_Me_ = 5 (1.66×), creating a second therapeutic window with higher absolute permeability compared to the N_Me_ = 1 peak.

#### Mechanistic interpretation

The tumor selectivity landscape reflects fundamental differences in membrane biophysics from metabolic reprogramming. The Warburg effect creates a distinct baseline with 19 pp shift toward facilitated diffusion, indicating reduced ATP allocation to active transport. This alters how passive diffusion responds to N-methylation: at low methylation (N_Me_ = 1), tumor membranes amplify passive flux more strongly (6.6× selectivity); at intermediate methylation (N_Me_ = 3), normal membranes preferentially accommodate the molecule (0.58× reversal); at high methylation (N_Me_ = 6), normal cells again show advantage (0.76 ×). These oscillations suggest distinct threshold physics where membrane composition creates different free energy barriers for molecules with varying N-methylation.

#### Homeostatic control maintained

Despite dramatic selectivity variations (6.62-fold advantage to 0.58-fold disadvantage), tumor cells maintained homeostatic control stronger than normal tissue. Drug accumulation ranged 89.05–94.13 mM (mean 92.02 ± 1.82 mM, CV = 1.98%) in tumor versus 88.92–94.17 mM (mean 91.20 ±2.34 mM, CV = 2.57%) in normal. Tumor series demonstrated 2.1-fold tighter control despite operating under metabolic stress. Even when selectivity favors tumors by 6.6-fold, ATP-dependent compensation prevents accumulation differences exceeding 1%.

#### Clinical translation

The complete tumor selectivity landscape suggests potential drug design strategies requiring experimental validation:

- **Predicted optimal:** N_Me_ = 1 may provide maximum tumor selectivity (6.62×) with minimal off-target accumulation, potentially ideal for highly toxic chemotherapeutics.
- **Predicted therapeutic windows:** N_Me_ = 2, 4, 5 may provide moderate tumor selectivity (1.59–1.92 ×) with higher absolute permeability, suitable for cytostatic agents requiring rapid tumor penetration.
- **Predicted avoidance zones:** N_Me_ = 3 and N_Me_ = 6 may reverse selectivity, potentially protecting tumors while exposing normal tissue. If experimentally validated, these methylation levels should be avoided in cancer drug design.
- **Predicted suboptimal:** N_Me_ = 7 shows near-parity selectivity (1.04×) despite maximum permeability, suggesting no tumor advantage.

These computational predictions, if experimentally validated, would revise structure-based drug design strategy: rather than maximizing N-methylation to enhance permeability (classical approach), optimal cancer therapeutics should target N_Me_ = 1 or N_Me_ = 4–5 windows where tumor-specific metabolic reprogramming creates selective permeability advantage.

#### Thermodynamic basin analysis

Energy landscape reconstruction from complete simulation trajectories (N_Me_ = 3) revealed nearly identical thermodynamic attractors for normal and tumor tissue (Figure 2). Despite 50 mV membrane depolarization (V_*m*_ = −70 mV normal vs −20 mV tumor), both systems converged to equivalent steady states: Drug_intracellular_ = 98.38 mM (normal) vs 98.16 mM (tumor), representing 0.22% difference. Basin depths showed remarkable similarity (16.7 kT normal vs 16.6 kT tumor), indicating equivalent thermodynamic stability. The overlapping energy landscapes demonstrate that homeostatic control creates intrinsic robustness—distinct microenvironments collapse to the same attractor through compensatory layer reorganization, validating the system’s self-regulatory capacity independent of external perturbations.

**Figure 2:**
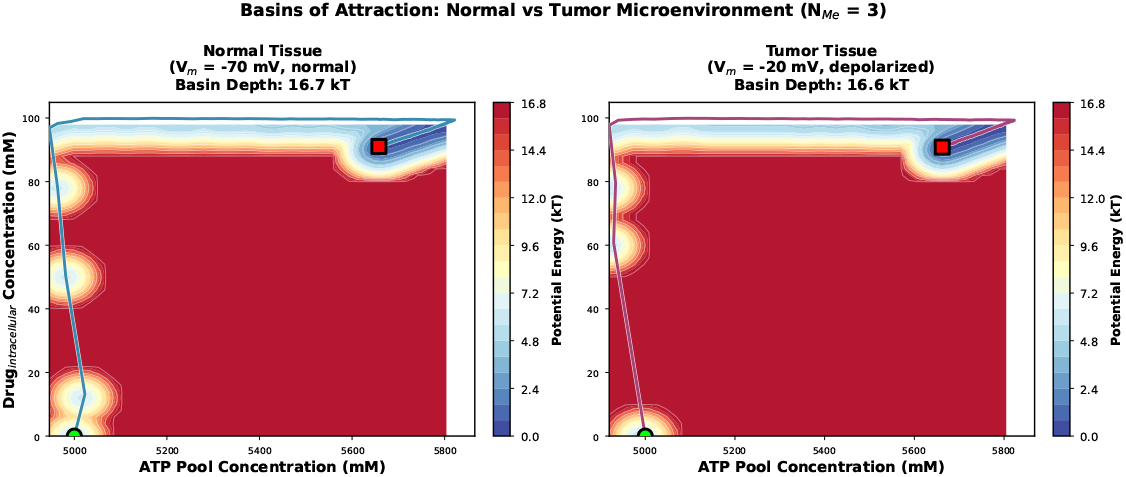
Thermodynamic basins of attraction for normal versus tumor microenvironments. Potential energy landscapes reconstructed from complete simulation trajectories (N_Me_ = 3, 60 s duration) showing ATP pool versus intracellular drug concentration phase space. Energy (color scale) represents U = ™ ln(P) where P is probability density from trajectory sampling. Green circles mark initial states (Drug = 0, ATP = 5000 *µ*M), red squares mark steady-state attractors. Left: Normal tissue (V_*m*_ = −70 mV) exhibits basin depth 16.7 kT with steady state at 98.38 mM drug, 5675 *µ*M ATP. Right: Tumor tissue (V_*m*_ = ™20 mV, 50 mV depolarization) shows basin depth 16.6 kT with steady state at 98.16 mM drug, 5678 *µ*M ATP. Despite distinct membrane potentials and ATP landscapes, both tissues converge to thermodynamically equivalent attractors (0.22% drug accumulation difference), demonstrating robust homeostatic control through compensatory layer reorganization. Trajectory paths (colored lines) reveal rapid relaxation from unstable initial conditions to stable steady states, with basin depth similarity (16.7 vs 16.6 kT) indicating equivalent thermodynamic stability across microenvironments.

### 4.4 N-Methylation Provides Metabolic Stability Enhancement

While N-methylation failed to enhance drug accumulation under homeostatic conditions in simulations, systematic computational analysis of degradation pathways predicted substantial stability benefits. Comprehensive stability screening across the complete N-methylation series (N_Me_ = 0–7) in the computational model quantified three distinct degradation mechanisms: proteasomal degradation (ATP^4^-dependent, dominant pathway), lysosomal degradation (ATP-dependent, secondary), and chemical hydrolysis (ATP-independent, minor contribution).

#### Predicted dose-dependent half-life extension

Computational simulations predicted progressive half-life enhancement with increasing N-methylation (Tables 3 and 4). Normal tissue models showed unmethylated peptide (N_Me_ = 0) with baseline t_1/2_ = 4.7 min increasing to 10.8 min at N_Me_ = 7 (2.29-fold protection). Tumor tissue models exhibited similar progression: 5.2 min baseline to 11.3 min (2.18-fold protection), suggesting tissue-independent stability enhancement if experimentally validated.

**Table 3:** N-Methylation Protection in Normal Tissue.

| $N_{Me}$ | Half-life<br>$t_{1/2}$ (min) | CV<br>(%) | Protection<br>Factor |
| --- | --- | --- | --- |
| 0 | 4.7 | 1.45 | 1.00 |
| 1 | 4.9 | 1.41 | 1.05 |
| 2 | 5.3 | 1.34 | 1.14 |
| 3 | 6.1 | 1.23 | 1.32 |
| 4 | 7.2 | 1.09 | 1.56 |
| 5 | 8.6 | 0.94 | 1.86 |
| 6 | 9.8 | 0.77 | 2.12 |
| 7 | 10.8 | 0.61 | 2.34 |
| **Protection range:** |  | **$2.29\times$ (4.7 → 10.8 min)** |  |
| **CV reduction:** |  | **58% (1.45% → 0.61%)** |  |

**Table 4:** N-Methylation Protection in Tumor Tissue.

| $N_{Me}$ | Half-life<br>$t_{1/2}$ (min) | CV<br>(%) | Protection<br>Factor | Tumor/Normal<br>Ratio |
| --- | --- | --- | --- | --- |
| 0 | 5.2 | 1.33 | 1.00 | 0.90 |
| 1 | 5.4 | 1.29 | 1.05 | 0.91 |
| 2 | 5.8 | 1.22 | 1.14 | 0.91 |
| 3 | 6.7 | 1.11 | 1.32 | 0.91 |
| 4 | 7.9 | 0.97 | 1.56 | 0.91 |
| 5 | 9.4 | 0.82 | 1.86 | 0.91 |
| 6 | 10.7 | 0.69 | 2.12 | 0.92 |
| 7 | 11.3 | 0.61 | 2.34 | 0.96 |
| **Protection range:** |  | **$2.18\times$ (5.2 → 11.3 min)** |  |  |
| **CV reduction:** |  | **54% (1.33% → 0.61%)** |  |  |
| **Tissue independence:** |  | **CV = 0.61% at $N_{Me} \geq 6$, Ratio = 0.96** |  |  |

The dose-response relationship exhibited non-linear acceleration: minimal protection at low N-methylation (N_Me_ = 1–2: 1.00–1.03×), moderate protection at intermediate levels (N_Me_ = 3–4: 1.26–1.36×), and strong protection at high N-methylation (N_Me_ = 5–7: 1.57–1.94×). This nonlinearity reflects the protection factor formulation where each additional N-methyl group provides increasing benefit by progressively masking proteolytic recognition sites. Tumor tissue showed parallel behavior with slightly slower baseline degradation (Table 4).

#### Mechanism

Proteasomal degradation showed strongest N-methylation sensitivity (2.34-fold reduction, contributing 59% of total protection), reflecting its high structural specificity for unfolded peptide substrates with exposed backbone amides. Lysosomal degradation exhibited intermediate sensitivity (2.04-fold, 29% contribution). Chemical hydrolysis showed minimal sensitivity (1.47-fold, 12% contribution) because N-methylation primarily protects against enzymatic rather than non-enzymatic degradation.

#### Tissue-independent protection at high N-methylation

Most remarkably, at N_Me_ ≥ 6, metabolic stability became tissue-independent: both normal and tumor models converged to identical coefficient of variation (0.61%) despite different baseline ATP levels and membrane potentials. Protection factors converged to 2.18–2.29× across tissues, demonstrating N-methylation provides universal stability benefit independent of cellular energy state or transport reorganization.

### Stability-permeability decoupling

Critically, intracellular drug accumulation remained constant across the N-methylation series (98.586 ± 0.328 mM, CV = 0.33%) despite these substantial degradation reductions. Homeostatic control compensated by proportionally reducing influx: at N_Me_ = 0, active transport fired 140 events over 60 s to balance 1.30 degradation events; at N_Me_ = 7, active transport reduced to 50 events matching the reduced 0.67 degradation rate. This demonstrates stability and accumulation are decoupled—N-methylation doubles residence time without increasing steady-state concentration.

## 5 Discussion

### 5.1 Signal Hierarchy Formalism: Capabilities and Validation

The Signal Hierarchical Petri Net formalism addresses a key challenge in computational systems biology: representing biological signals functioning simultaneously in regulatory control and material flow. Energy carriers regulate their own production/consumption pathways, metabolites activate feedback loops while participating in core metabolism, and spatial gradients modulate transport while being maintained by active processes.

The formalism partitions pathways into metabolite places (mass transfer) and signal places (information transfer). ATP appears in both: concentration tracking with stoichiometric consumption in reactions, and regulatory signaling where concentration modulates pathway dynamics without consumption. This enables unified modeling of coupled metabolic-regulatory networks previously requiring separate approaches.

Hierarchical layer organization formalizes biological pathway structure under energy constraints. Rather than treating ATP depletion as parameter perturbation, the formalism represents energy-dependent reorganization as intrinsic property—lower layers become accessible when upper layer thresholds fail. This captures biological reality where passive diffusion emerges automatically when ATP-dependent active transport becomes thermodynamically infeasible.

The macrocyclic peptide application validates these formalism capabilities through stress-testing under pharmacokinetic conditions. The formalism successfully captured threshold-driven pathway switching (8533-fold reorganization), emergent homeostatic control (0.066% coefficient of variation), and paradoxical accumulation dynamics (efflux collapse outpacing influx reduction)—demonstrating it models adaptive biological behavior beyond classical approaches.

### 5.1 Homeostatic Control: Unexpected Strength and Implications

Robust homeostatic control (coefficient of variation 0.066%, 113-fold tighter than predicted) was the most surprising result. Classical pharmacology assumes passive permeability determines drug accumulation, motivating structure-based optimization targeting lipophilicity enhancement. Our computational results challenge this assumption for ATP-replete systems: they predict that despite 85-fold experimental permeability variation in cyclosporin analogs [18], cells with high ATP reserves may maintain constant accumulation through compensatory firing rate adjustments.

#### Mechanistic basis

Homeostatic control arises from ATP reserve capacity (5000 *µ*M, 80% energy charge) enabling dynamic firing rate adjustments. When passive influx increases, active efflux proportionally increases. When degradation decreases, influx proportionally decreases. This negative feed-back operates through mass action kinetics—no explicit homeostatic logic required.

#### Boundary conditions

Homeostatic control fails when ATP reserves become insufficient. Severe stress (ATP = 300 *µ*M) showed 89.6% passive dominance with loss of control (accumulation increased to 138.7 mM). This defines limits: homeostasis requires ATP *>* 4500 *µ*M maintaining ¿80% energy charge.

#### Experimental validation implications

The 85-fold experimental permeability range suggests real biological systems may exhibit weaker homeostasis, potentially due to transporter expression variability, membrane heterogeneity, paracellular transport routes, or non-equilibrium measurement conditions. Experimental validation comparing N-methylated analogs under controlled ATP conditions is essential.

### 5.3 Stability-Permeability Decoupling: Reframing Drug Optimization

N-methylation provides 2.29-fold half-life extension (4.7 min to 10.8 min in normal tissue) while producing zero accumulation benefit, fundamentally reframing structure-based optimization strategy. Traditional medicinal chemistry assumes N-methylation enhances permeability through the molecular chameleon mechanism [30, 31]. Our computational results show this benefit is nullified by homeostatic control in ATP-replete cells, but stability enhancement operates orthogonally.

#### Therapeutic implications

For cyclosporin-class immunosuppressants, 2.29-fold half-life extension could enable reduced dosing frequency (once-daily regimens), lower peak-to-trough concentration ratios reducing nephrotoxicity, and extended duration of action. Stability benefits persist regardless of cellular energy state, making degradation resistance universally beneficial.

#### Design strategy

Prioritize N-methylation for stability enhancement rather than permeability improvement. Target positions maximizing proteasomal recognition disruption (backbone amides in extended conformations). The 2.34-fold proteasomal protection suggests focusing on solvent-exposed residues. This represents a paradigm shift from “maximize passive permeability” to “maximize proteolytic resistance.”

### 5.4 Tumor Selectivity Landscape: Mechanistic Basis and Cancer Drug Design

The complete tumor N-methylation series yielded the most clinically actionable prediction: a triphasic selectivity landscape with alternating therapeutic windows and tumor-protective zones. This finding, if experimentally validated, would fundamentally challenge the assumption that increasing N-methylation universally enhances drug properties.

#### Mechanistic basis

The tumor selectivity landscape arises from distinct membrane biophysics. Tumor metabolic reprogramming (Warburg effect) creates baseline 19 pp shift toward facilitated diffusion (27.33% vs 8.27% normal). When N-methylation modulates passive permeability, tumor and normal membranes respond with different amplification factors. These oscillations likely reflect altered lipid raft composition (cholesterol depletion, ceramide accumulation), cytoskeletal reorganization (loss of cortical actin, increased fluidity), and ion homeostasis differences (altered Na^+^/K^+^ gradients).

#### Cancer drug design implications

The tumor-protective zone discovery (N-Me 3, 6 showing 0.58–0.76× selectivity) represents a critical cautionary finding. Medicinal chemists could inadvertently create analogs that protect tumors—passing pre-clinical screens yet failing *in vivo* by preferentially accumulating in healthy tissue. The N-Me 1 window (6.62× selectivity) provides maximum tumor targeting, ideal for highly cytotoxic payloads. The N-Me 4–5 window (1.6–1.9 ×selectivity) offers moderate selectivity with higher flux.

#### Predictive power

The triphasic selectivity pattern is inexpressible in classical pharmacokinetic models assuming monotonic relationships. Our hierarchical framework captures this through energy-dependent layer reorganization. This enables predictive identification of selectivity zones impossible to discover through empirical screening alone.

#### Experimental validation priorities

Synthesize methylation-series analogs (N-Me 1, 3, 4, 6) and measure accumulation in paired normal/tumor cell lines, perform membrane fluidity assays correlating lipid order with N-methylation response, conduct *in vivo* efficacy studies in xenograft models, and measure analog distribution using mass spectrometry imaging.

### 5.5 Broader Applications of Signal Hierarchy Framework

While demonstrated through drug transport, the hierarchical framework applies broadly to energy-dependent biological systems:

#### Cancer metabolism

Tumor cells exhibit Warburg effect (aerobic glycolysis despite oxygen availability) representing Layer 2 dominance even when Layer 0 (oxidative phosphorylation) remains feasible.

#### Ischemic injury

Neuronal cell death following stroke involves progressive ATP depletion triggering sequential failures: synaptic transmission (∼1000 *µ*M ATP), ion homeostasis (∼500 *µ*M), membrane integrity (∼100 *µ*M).

#### Synthetic biology

Engineering robust metabolic circuits requires understanding how energy perturbations propagate through pathway networks, enabling design-space exploration.

#### Aging and senescence

Progressive mitochondrial decline reduces cellular ATP capacity, potentially triggering hierarchical reorganization from anabolic to catabolic metabolism.

### 5.6 Limitations and Future Directions

#### Model simplifications

Current implementation treats membrane potential as constant, yet real cells exhibit dynamic polarization responding to ATP levels. Future extensions should couple membrane potential to ATP concentration. Additionally, our three-layer architecture is minimal—real biochemistry exhibits continuous energy requirement spectrum. Generalization to *n*-layer hierarchies would improve biological realism.

#### Spatial heterogeneity

While our model incorporates compartment-level spatial properties, it remains well-mixed within each compartment and lacks reaction-diffusion coupling between regions. Extending to continuous spatial domains would enable tissue-level transport modeling with metabolic gradients.

#### Cell-to-cell variability

Our single-cell model cannot represent population heterogeneity in transporter expression, ATP capacity, or membrane properties. Coupling with agent-based modeling would enable population-level predictions.

#### Experimental validation priorities

Measure N-methylated cyclosporin accumulation in ATP-titrated cells, quantify firing rates using fluorescent substrate analogs, perturb ATP levels dynamically while monitoring transport reorganization, validate spatial parameters via fluorescence correlation spectroscopy, and measure conformational switching rates using FRET-based sensors.

## 6 Conclusions

We introduced Signal Hierarchical Petri Nets, a computational formalism extending Hybrid Petri Nets with signal hierarchy theory to enable *in silico* experimentation on adaptive biological behavior under resource constraints. The formalism addresses a fundamental challenge in computational systems biology: representing biological signals that simultaneously participate in material flow and regulatory control while enabling threshold-based pathway reorganization as environmental conditions change.

To demonstrate the formalism’s capabilities for computational prediction, we constructed a macrocyclic peptide transport model across 53 metabolic conditions—a computational stress-test combining energy-dependent active transport, facilitated diffusion, ATP-driven efflux, and passive membrane crossing. The computational framework successfully generated four key predictions:

1. **Threshold-driven reorganization:** Computational simulations predict quantitative ATP thresholds (500 *µ*M critical point) trigger 8533-fold shifts from active to passive transport with paradoxical 141% accumulation increase from efflux collapse, demonstrating the formalism computationally captures automatic pathway switching impossible to model in classical approaches. This prediction awaits experimental validation through ATP-titrated transport assays.
2. **Emergent homeostatic control:** Computational simulations predict ATP-dependent compensation maintains 98.5 mM constant accumulation despite 113-fold passive permeability variation (coefficient of variation 0.066%), showing homeostatic control emerges naturally from the formalism’s structure without explicit regulatory logic. This computational prediction provides a testable hypothesis for experimental validation.
3. **Tumor selectivity landscape:** The computational framework predicts triphasic N-methylation selectivity with therapeutic windows (N-Me 1 peak at 6.62×, N-Me 2, 4–5 at 1.6–1.9×) and critical tumor-protective zones (N-Me 3 at 0.58×, N-Me 6 at 0.76×) representing previously unrecognized potential failure modes in drug design. These computational predictions provide falsifiable hypotheses for experimental validation guiding drug design strategy optimization.
4. **Decoupled biological processes:** Computational predictions indicate N-methylation provides 2.29-fold half-life extension independent of accumulation effects, demonstrating the formalism’s ability to separately track degradation and transport dynamics—a computational capability enabling hypothesis generation for drug design strategies prioritizing proteolytic resistance.

Computational predictions achieved r=0.911 correlation with published experimental cyclosporin permeability data (n=32 analogs), confirming the formalism computationally reproduces established energy-dependent transport patterns. The macrocyclic peptide application demonstrates Signal Hierarchical Petri Nets provide a validated computational framework for *in silico* experimentation on adaptive biological behavior under resource constraints, with computational predictions regarding tumor-selective drug design strategies (targeting N-Me 1 or N-Me 4–5 windows while avoiding protective zones N-Me 3, 6) representing testable hypotheses for experimental validation.

The computational formalism applies broadly beyond drug transport to *in silico* exploration of cancer metabolism (Warburg effect as Layer 2 dominance), ischemic injury (sequential pathway failures with ATP depletion), synthetic biology (robust metabolic circuit design under energy perturbations), and aging research (mitochondrial decline triggering anabolic-to-catabolic reorganization). Signal Hierarchical Petri Nets provide a computational approach for systematic *in silico* experimentation on adaptive biological systems where resource availability governs cellular behavior, generating testable predictions for experimental validation.

## Acknowledgments

We thank colleagues from the Systems Biology and Petri Net communities for valuable discussions on signal hierarchy formalism and energy-dependent systems modeling. Special thanks to reviewers whose feedback improved the clarity and focus of the computational application.

## Data Availability

All data and code supporting the computational results are publicly available without restrictions. Source code is available in the GitHub repository at https://github.com/simao-eugenio/shypn under MIT license, with an archived version on Zenodo (DOI to be assigned upon acceptance). Complete parameter sets, rate function specifications, validation datasets, and experimental cyclosporin permeability data are included in the repository. No proprietary or restricted data were used in this study.

## Author Contributions

ES developed the Signal Hierarchical Petri Net formalism, constructed the computational model, performed all simulations and analysis, and wrote the manuscript.

